# A systematic assessment of machine learning for structural variant filtering

**DOI:** 10.64898/2026.01.27.702059

**Authors:** Archit Kalra, Luis F Paulin, Fritz J Sedlazeck

## Abstract

**Background:** Accurate discrimination of true structural variants (SVs) from artifacts in long-read sequencing data remains a critical bottleneck. Numerous machine learning solutions have been proposed, ranging from classical models using engineered features to advanced deep learning and foundation model interpretability methods. However, a systematic comparison of their performance, efficiency, and practical utility is lacking.

**Results:** We conducted a comprehensive benchmark of five machine learning paradigms for SV filtering using standardized Genome in a Bottle (GIAB) data for samples HG002 and HG005. We evaluated classical Random Forest classifiers on 15 genomic features, computer vision models (ResNet/VICReg), diffusion-based anomaly detection, sparse autoencoders (SAEs) on the Evo2-7B foundation model, and multimodal ensembles. A simple Random Forest on interpretable features achieved a peak F1-score of 95.7%, effectively matching all more complex models (ResNet50: 95.9%, Diffusion: 95.8%). This study represents the first application of diffusion-based anomaly detection and sparse autoencoders to structural variant analysis; while diffusion models learned highly discriminative, disentangled representations and SAEs uncovered biologically interpretable features (including atoms that were specific for ALU deletions, chromosome X variants and insertion events), they did not significantly surpass this classification ceiling. Ensemble methods offered no performance benefit but may have future potential given the orthogonality of vision-based and linear features.

**Conclusions:** Our findings demonstrate that for the established task of germline SV filtering, simpler, interpretable models provide an optimal balance of accuracy, speed, and transparency. This benchmark establishes a pragmatic framework for method selection and argues that increased model complexity must be justified by clear, unmet biological needs rather than marginal predictive gains.

## Introduction

Structural Variant (SV) detection is becoming more mature over the past years thanks to multifold developments on algorithmics, improvements in sequencing accuracy and availability of benchmark data sets [1,2]. This leads to multiple successes on the identification and interpretation of SV impact across diseases such as neurological, cardiovascular diseases and even cancer [3–5]. While long-reads provide key benefits they also have their own limitations that come from ascertainments of high complexity regions, higher error rates in repeats and chimeric artifacts impacting germline and somatic SV identification[6–9]. This still leads to some false discovery rates that are becoming more obvious once comparing samples eg. to detect de novo variants in a family as an example or somatic SV [9,10].

To improve upon this multiple filtering approaches have been implemented most often within the SV calling methods themselves. For example, Sniffles[6] has multiple filters that are deployed based on coverage, read support, breakpoint accuracy to name only a few features. These heuristics, while computationally efficient, often are somewhat ad hoc and might not adequately work in multiple scenarios. For this and other reasons genotype methods such as Kanpig[10] or SVJedi [11] are still popular to provide independent quantification of presence or absences of SV [12]. Still challenges remain to provide dynamic and most accurate filters that can more rapidly adapt to sequence content, sequencing technologies, coverage fluctuations and complexities of the sample (e.g. ploidy).

Over the past few years multiple Machine learning (ML) methods have been introduced and adopted to address some of the needs. This has been especially successful in the detection and filtering of SNV and indel candidates[13–15]. A few methods have been proposed over the past years that leverage some sort of ML to improve SV detection; DeepSVFilter [16] was among the first to apply deep learning to SV filtering, using convolutional neural network (CNN) and recurrent neural network (RNN) models to classify variants based on alignment features encoded as images. While it demonstrated that machine learning could learn to distinguish real from spurious SV signals, DeepSVFilter was limited to short-read data and occasionally eliminated true variants along with FPs. Building on this foundation, [17] AquilaDeepFilter was built for short reads and linked reads, while Xia et al. [18] developed CSV-Filter, which introduced a more sophisticated multi-channel image encoding of local alignments based on CIGAR strings. By fine-tuning residual neural network (ResNet) models and using variance-invariance-covariance regularization (VICReg) to prevent model collapse, CSV-Filter improved precision by approximately 15% for various callers without substantially reducing true positive (TP) detection. Thus, visual representations of alignment evidence, encoding coverage patterns, CIGAR operations, and local sequence context, have been shown to help boost variant caller precision [18]. However, the success for filtering SVs using ML has been modest, to say the least, with not many tools leveraging this emerging trend. Indeed, this might be because of limiting training data although more and more well curated genomic SV data sets are emerging for example Genome in a Bottle (GIAB), Human Pangenome Reference Consortium (HPRC) and many others [19,20]. Simultaneously there have been many new approaches in ML introduced to tackle the problem of SV ascertainment. Initial efforts employed classical models on curated genomic features, followed by more complex deep learning architectures that operate directly on raw alignment data (e.g., image-based convolutional networks). Most recently, generative models and foundation language models have entered the arena, promising not only improved accuracy but also novel avenues for interpretability. Despite this rapid methodological expansion, a systematic and equitable comparison across these fundamentally different paradigms is lacking, leaving scientists without clear guidance on which approach to deploy.

In this study we benchmark five distinct ML strategies for SV filtering: (1) classifiers using engineered genomic features, (2) computer vision models trained on alignment images, (3) diffusion-based anomaly detection, (4) sparse autoencoders applied to a genomic foundation model (Evo2-7B), and (5) multimodal ensemble architectures. Using standardized Genome in a Bottle (GIAB) benchmarks for samples HG002 and HG005, we evaluate each method in terms of precision, recall, computational efficiency, and interpretability. This work represents the first comprehensive cross-paradigm evaluation of ML approaches for SV filtering, including novel applications of diffusion-based anomaly detection and sparse autoencoder interpretability methods to genomic variant analysis. Our goal is to establish a clear performance landscape and to determine whether increased model complexity translates into tangible gains for real-world SV filtration tasks. As such this work provides deeper insights into the individual models currently available and their potential benefits and shortcomings.

## Results

### A comprehensive benchmark for machine learning-based SV filtering

To establish a rigorous comparative framework, we evaluated five machine learning paradigms for filtering false positive SVs from 25x or higher coverage Oxford Nanopore long-read SV call sets (**Figure 1A**). Our benchmark is built on the GIAB high-confidence SV sets for samples HG002 and HG005, aligned to both GRCh37 and GRCh38 references [21,22]. The final dataset comprised 50,795 SVs (44,842 TPs, 5,953 FPs) composed of 50bp or larger insertion and deletions from HG002 and HG005, partitioned across three genome-reference combinations to enable both within- and cross-sample evaluation (**Figure 1B**, see Methods for details). We benchmarked: (1) traditional classifiers on 15 engineered genomic features, (2) computer vision models (ResNet, VICReg) trained on 4-channel alignment images, (3) diffusion-based anomaly detection models, (4) interpretable sparse autoencoders (SAEs) applied to the Evo2-7B genome foundation model, and (5) multimodal ensemble architectures. All models were evaluated against Sniffles2[6] call sets using Truvari [23], with performance measured by precision, recall, and F1-score (**Table 1**).

**Figure 1:**
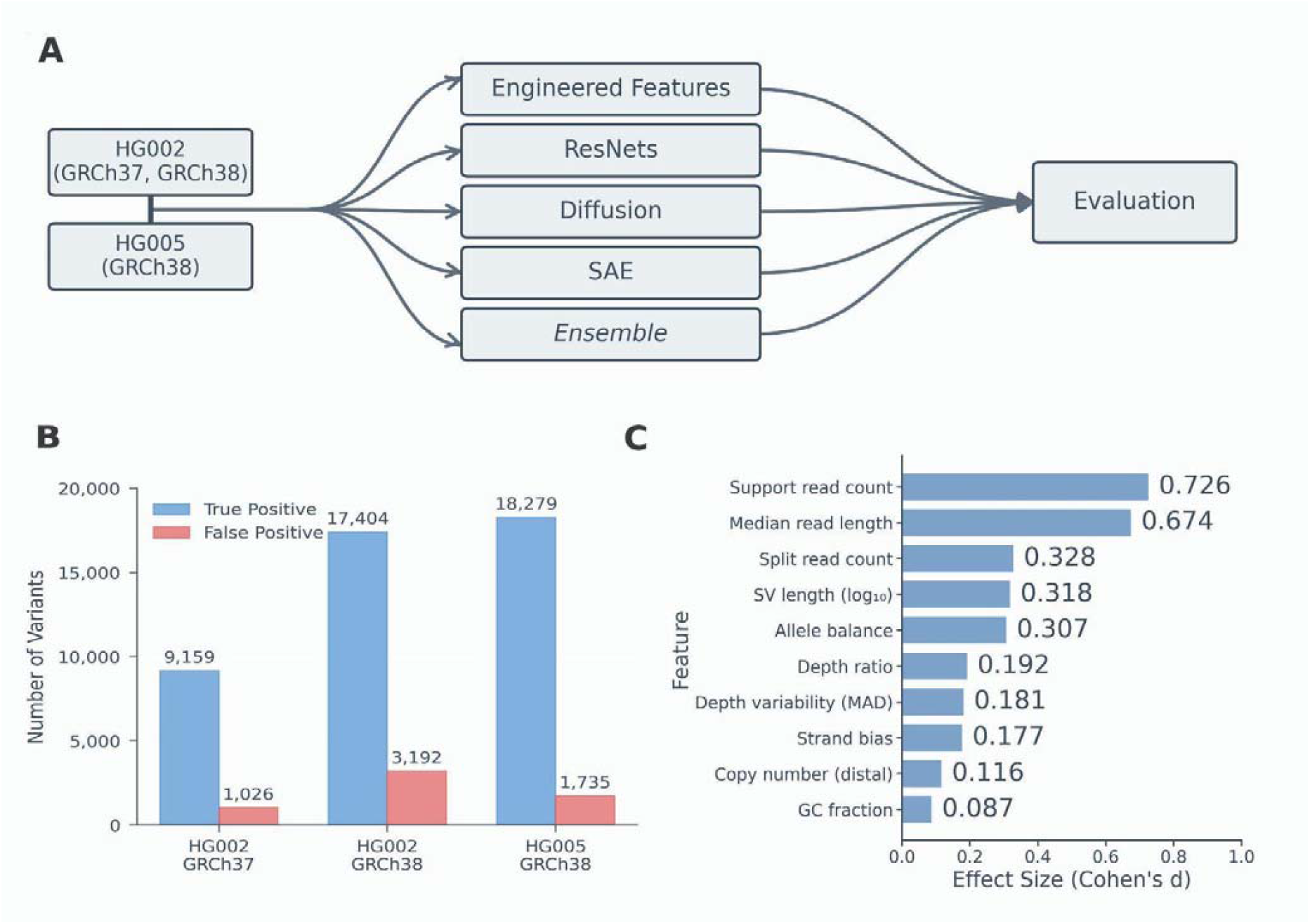
Study Framework & Core Performance. A. Benchmark workflow: The benchmark datasets for HG002 with GRCh37 and GRCh38 alignments, and HG005 with the GRCh38 alignment are used to train models based on Engineered linear features, Vision, Diffusion, SAE and Ensemble structures with linear and vision-based features. Models are trained and evaluated using an 80/20 split of the data described in (B). B. The data composition of the study included insertions and deletions from the HG002_GRCh37, HG002_GRCh38, HG005_GRCh38 datasets, with Sniffles2 calls compared to the Genome In A Bottle benchmark to establish true positives and false positives. This led to a total of 50,795 SV for training and evaluation (HG002_GRCh37: 9,159 TP; HG002_GRCh38: 17,404 TP; HG005_GRCh38: 18,279 TP). C. Engineered feature importance: Cohen’s d effect sizes for top 10 features in terms of statistically significant (p < 0.01) differences in distribution across all three datasets. Support read count (0.726) and median read length (0.674) are strongest discriminators between true and false positive SVs. Genomic context features (GC fraction, copy number) show minimal effect.

We first investigated the sufficiency of hand-engineered genomic features for discriminative filtering. We defined 15 features spanning four functional categories: number of support reads, median read length, SV length, allele balance, number split reads, strand bias, depth mean absolute deviation, depth ratio, ratio of coverage in the SV vs flanking ends and GC fraction (**Table 2**). Mann-Whitney U tests and Cohen’s *d* were used to evaluate differences between the distributions for TP and FP for each feature for SVs across HG002 and HG005. 13 of the 15 features exhibited significant (p < 0.01) separation between TPs and FPs in all benchmark datasets. The most consistently discriminative features were number of support reads (effect size d ≥ 0.59 across datasets), depth ratio, and log of SV length, showing the fundamental role of read support and local coverage imbalance in SV validation (**Figure 1C**).

A Random Forest (RF) classifier trained on all 15 features achieved a peak F1-score of 95.7%, representing a 2-percentage-point improvement over the Sniffles2 baseline (93.8% F1) (**Table 1**). This classifier demonstrated an optimal balance, filtering out 46.8% of FPs (558/1,191) while discarding only 1.6% of TPs (143/8,968) on the held-out test set. The RF significantly outperformed logistic regression on the same features (94.4% F1), indicating that non-linear interactions, such as between read support and mapping quality drop, carry additional discriminative power. Feature ablation studies confirmed robustness: models using only the top 8 or 10 features retained F1-scores above 94.9%, while PCA-reduced features performed slightly worse (94.9% F1). This establishes a strong, interpretable benchmark where each predictive component maps directly to a measurable genomic or technical property.

### Deep learning models achieve parity in accuracy but diverge in representation and cost

We next assessed whether more complex, data-driven representations (see methods) could surpass this engineered-feature baseline. Given the success of CSV-Filter in using fine-tuned ResNet models, we evaluated multiple vision models on the same data sets to be able to compare their results.

Vision models, including ResNets and U-Nets, were trained on 224×224 pixel images encoding CIGAR operations (match, deletion, insertion, soft-clip), using an image generation technique previously outlined by Xia et al. [18] (see methods). A standard ImageNet-pretrained ResNet50 achieved an F1-score of 95.9%, statistically matching the RF benchmark. VICReg self-supervised models showed strong cross-genome generalization, with a VICReg-trained ResNet50×2 model maintaining 92.3% accuracy when trained on HG002 and tested on HG005. However, their internal representations proved less immediately useful: logistic regression on extracted ResNet50 latents yielded only 94.5% F1, and three-class classification (INS/DEL/FP) failed entirely (accuracy <60%, AUPRC=0.000).

Diffusion models were implemented as anomaly detectors, trained exclusively on TP samples to learn the “manifold” of true SV signatures. This is due to the expectation that FP SVs represent scattered technical artifacts rather than coherent biological patterns, an assumption validated by unsupervised clustering analysis of learned representations from initial model training runs. For each test sample, forward diffusion adds noise to the input image and the trained model predicts the added noise. The mean squared error between these is the reconstruction error, and used as a classification score. The lower the reconstruction error, the higher the TP likelihood ([24]; [25]). Through systematic architecture and noise-schedule sweeps (**Table 3**), we identified a U-Net Large model with a fast schedule (500 timesteps) as optimal. After moving the reconstruction-error threshold to maximize the F1 score, this model achieved an F1-score of 95.8%, with high recall (99.5%) and precision (92.5%) (**Figure 2A**). No significant differences were noted between the misclassified and correctly classified FPs in any of the 15 genomic features (all p > 0.05), suggesting diffusion models access information beyond standard linear representations.

**Figure 2:**
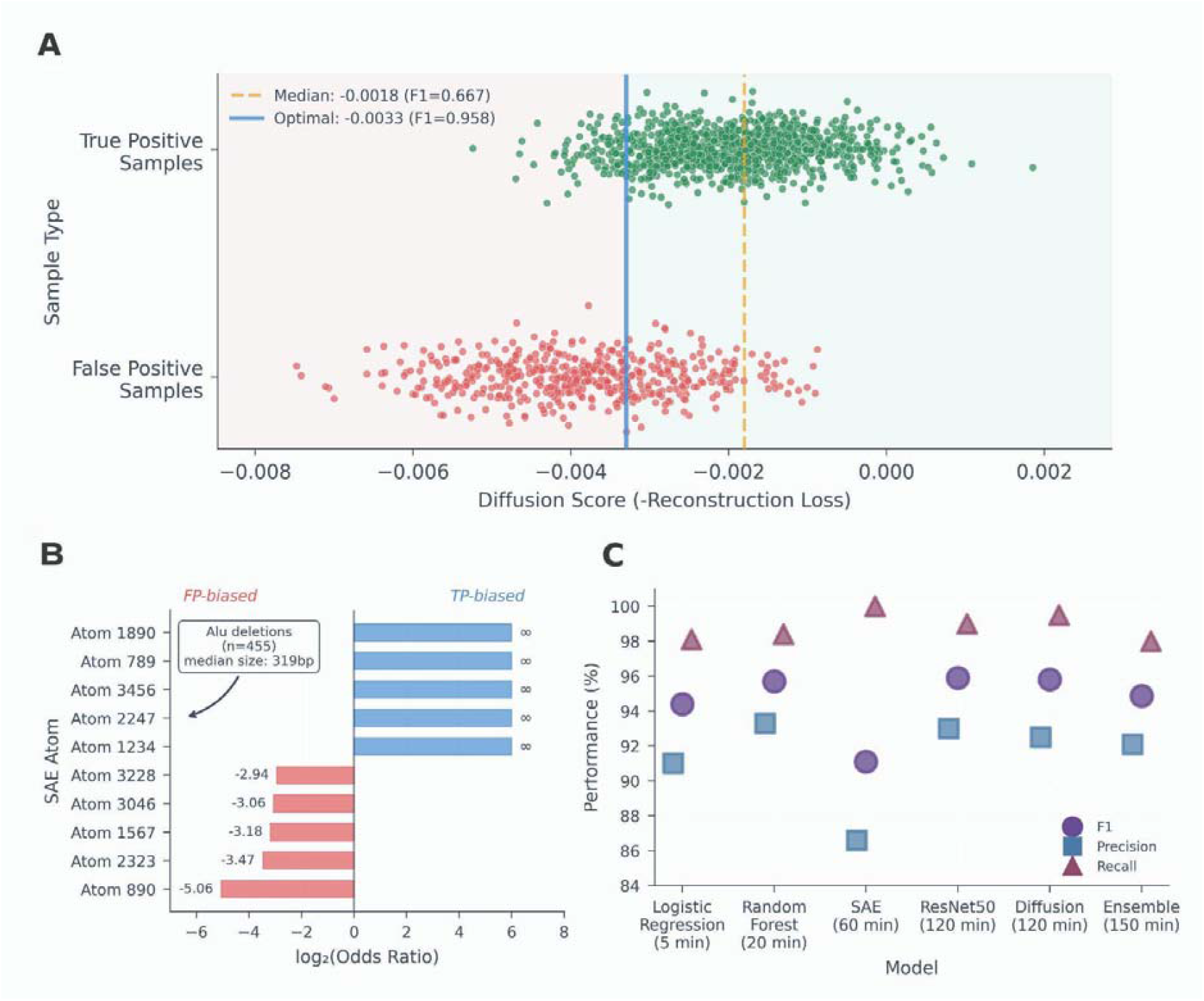
Model-Specific Analyses & Efficiency. A. Optimization of reconstruction error threshold for FP classification via diffusion model. Though the model did successfully filter out a significant number of FP by classifying all samples below the median as FP, this also eliminated nearly half of TP. Moving the reconstruction error clear of the majority of TP boosted the F1 score by 0.29 points. B. SAE atom specialization: Top 10 interpretable atoms by odds ratio. FP-biased atoms (red) may indicate technical artifacts of the variant caller. TP-biased atoms (blue) capture high-confidence structural variants. C. Performance-efficiency tradeoff: F1, precision, and recall versus training time. Diffusion (95.8% F1) and ResNet50 (95.9% F1) offer fine-tuning / training time on the order of a few hours for best performance. Random Forest (95.7% F1) trains in a fraction of the time and also offers a rapid baseline.

Sparse autoencoders (SAE) applied to the Evo2-7B foundation model offered a distinct advantage: interpretable feature discovery. Evo 2-7B, a genomic foundation model with 7B parameters and trained on 9.3 trillion prokaryotic and eukaryotic DNA base pairs with single-nucleotide resolution, was used to extract embeddings from 1000 bp windows surrounding SV breakpoints [26]. We trained 31 SAE configurations on the activations in Layer 26 of the Evo2 model (previously used by the AI interpretability firm Goodfire for analysis with strong results), identifying a 4,096-dimensional, Top64 sparsity SAE (SAE_4096_64) that balanced performance and efficiency ([27]. Of the 3,700 features with more than five activations on the test set, 812 exhibited a strong TP bias (odds ratio > 2.0) and 377 had a strong FP bias (odds ratio < 0.5). This model, 8 times smaller than the prior Goodfire SAE (trained on Evo2-7B activations on prokaryotic and eukaryotic sequences), achieved a peak hard-filter F1 (where SVs activating the most FP-biased atoms were classified as FP - see methods) of 91.1% by simply thresholding activations of FP-biased features. More interestingly, the odds ratio bias uncovered biologically meaningful “atoms,” which are summarized in **Supplementary Table 1**. For instance, Atom 2247 activated strongly on ALU-mediated deletions, with 98.7% of atom-activating deletions and 99.1% of Alu-sized variants (250-350bp) confirmed to overlap RepeatMasker[28] ALU annotations and a median SV size of 319 bp (**Figure 2B**). Other atoms specialized for insertions (e.g., Atom 3987, 100% INS bias), specific chromosomes (Atom 907, 98.4% on chrX), or technical artifacts (Atom 3228, 100% FP bias). Beyond general TP/FP differences, variants activating the ALU specialist, Atom 2247, showed elevated homopolymer content compared to other variants (+1.25 standard deviations, p=5.40e-15), indicating learned recognition of ALU-specific sequence signatures such as poly-A tails. Visualization via UMAP revealed TPs forming dense, structured clusters (75 clusters, median neighbor distance 0.135), whereas FPs were more diffusely scattered (12 clusters, median distance 0.310), corroborating the SAE’s ability to separate biologically coherent patterns from noise. Importantly, these feature associations formed without any supervision; the SAE atoms *learned* these patterns without any labeling as such.

### Multimodal integration reveals saturation of discriminative information

Despite their differing architectures, all models converged on a similar performance maximum. We reasoned that their underlying representations might capture orthogonal aspects of SV biology. Engineered features codify known technical biases, diffusion latents isolate novel visual signatures, and SAE atoms extract discrete genomic concepts. We systematically tested whether combining these orthogonal views could push classification accuracy beyond the limit achievable by any single method. **Figure 2C** describes the overall performance results for all evaluated models.

We first quantified redundancy via canonical correlation analysis (CCA). This confirmed high correlation (r ≥ 0.95) among the three vision-based modalities (diffusion, ResNet, VICReg latents) and separately among the scalar features (engineered features and SAE activations) (**Figure 3A**). Mutual information analysis was also conducted among latents extracted from the backbone of each model. Strikingly, an analysis of the 2,816 latents extracted from the U-Net’s bottleneck revealed them to be a uniquely disentangled representation: they exhibited the lowest mutual information (0.0006 bits) and the highest proportion of discriminative features (94.8%) among all tested modalities (**Figure 3B**). This suggests diffusion models learn a set of nearly independent, highly informative features, yet a Random Forest on these latents reached only 93.8% F1, indicating that the final classification step, not feature quality, becomes the limiting factor.

**Figure 3:**
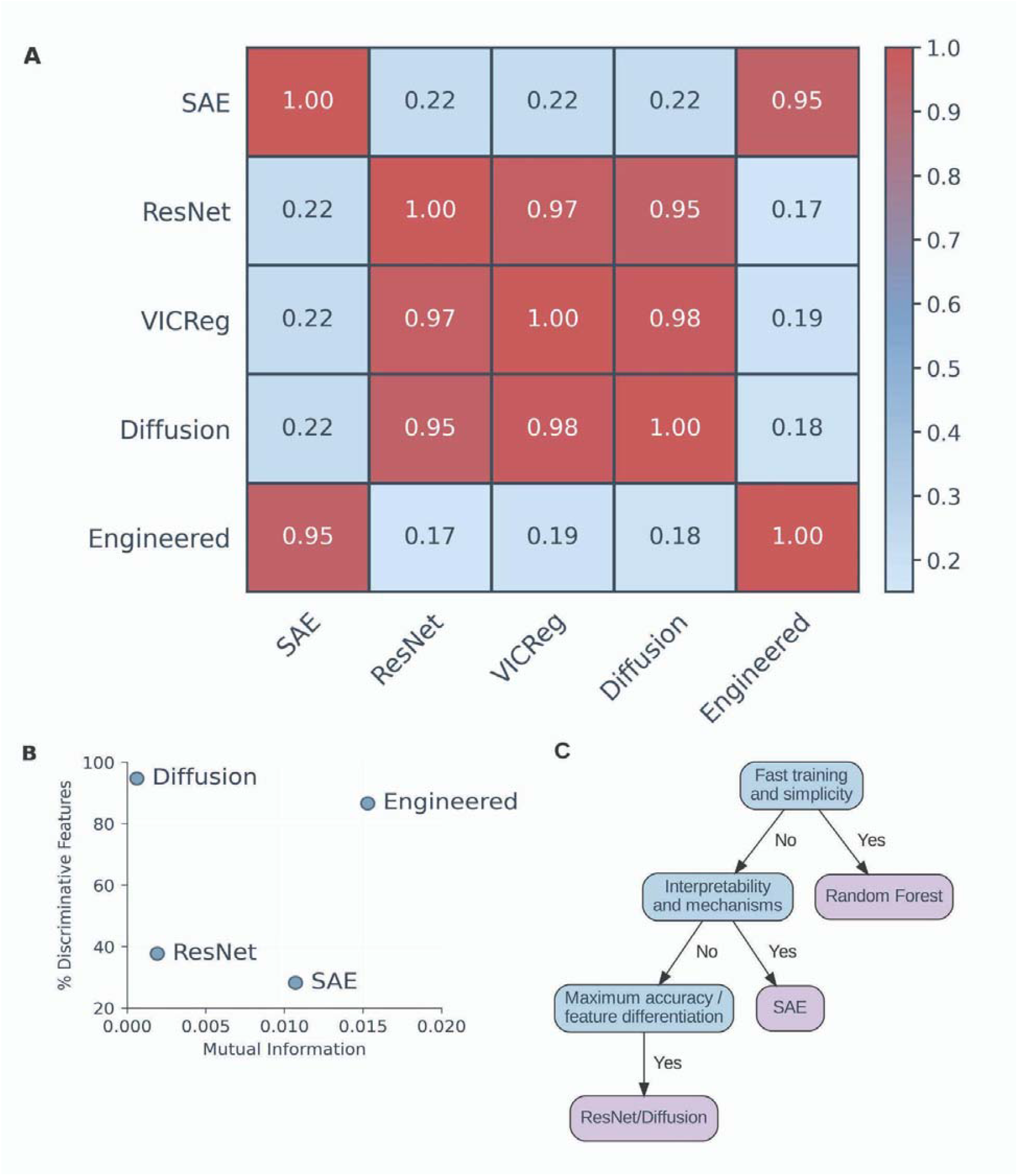
Feature Analysis & Practical Guidance. A.Cross-modal canonical correlation analysis (CCA) of five modalities used for SV filtering experiments. The image modalities (diffusion, ResNet, VICReg) had high correlations with each other, and the scalar modalities (linear, SAE) similarly had high correlations with each other. B.Mutual information versus discriminative feature percentage across representation types. Diffusion latents achieve the highest discriminative power (94.8%) with minimal mutual information (0.0006 bits), indicating independent, informative features. Engineered features show highest redundancy (0.0153 MI, 86.7% discriminative). C.Decision flowchart: Decision flowchart for practical applications. Random Forest is most useful for speed, high accuracy and simplicity, SAE for interpretability and mechanism discovery, and ResNet/Diffusion for maximum accuracy and downstream feature differentiation.

We subsequently developed an attention-based fusion network to integrate the most promising, non-redundant modalities: diffusion latents and engineered features. This ensemble achieved an F1-score of 94.9%, which is failing to surpass the best single-modality models (RF: 95.7%, Diffusion: 95.8%) (**Table 4**). Similarly, other ensemble strategies and neural network classifiers on SAE features (best SAE NN F1: 92.4%) could not close the gap to the simple RF benchmark.

This convergent result indicates that for the task of SV filtering on high-quality benchmarks, the discriminative signal is saturated at approximately 96% F1. The added complexity, computational cost (the large vision models took on the order of hours to train, as opposed to the linear models, which took on the order of 20-30 minutes), and loss of interpretability associated with deep learning and fusion approaches are therefore not justified by incremental gains (**Figure 3C**). Instead, the choice of method should be guided by secondary priorities: interpretability and speed favor engineered features with Random Forest; investigation of foundation model reasoning favors SAEs; and exploration of novel, disentangled representations favors diffusion models.

## Discussion

Our benchmark reveals a clear and perhaps unexpected result: for the core task of filtering false positive structural variants (SV) from long-read call sets, a simple Random Forest model trained on 15 interpretable genomic features defines a robust performance ceiling, achieving an F1-score of 95.7%. This result matches or exceeds the performance of all significantly more complex deep learning models we evaluated. This has actually multiple advantages such as that we can easier understand why certain models work based on expected features. Furthermore, these more simplistic models achieve their task in generally faster and with less compute requirements, thus retaining time for other improvements in SV calling such as localized assemblies, or general fast turn around times for variant calling.

This conclusion, however, does not diminish the substantial methodological contributions of advanced deep learning models, but instead refines their role. To our knowledge, this study represents the first application of diffusion models and sparse autoencoders to variant calling, demonstrating capabilities beyond classification. While they did not outperform as classifiers, they provided unique and complementary insights. Diffusion models, for instance, learned a uniquely disentangled latent representation, exhibiting the lowest mutual information and the highest proportion of discriminative features among all modalities. This suggests they excel not at final classification, but at distilling the complex visual signature of an SV into a set of nearly orthogonal components, which is a powerful property for exploratory analysis or feature engineering. Similarly, sparse autoencoders applied to the Evo2-7B foundation model functioned not as superior filters, but as powerful interpretability engines. They automatically isolated discrete, biologically meaningful concepts without any supervision. For example, several features are specific to Alu-mediated deletions or chromosome X SV directly from the model’s activations. Thus, the primary utility of these highly sophisticated approaches may shift from being the discovery tool to generate novel hypotheses on the impact of SV or even the mechanism of their occurrence. Notably, our benchmark employs a different negative sample strategy than prior studies, such as CSV-Filter. Xia et al. [18] generate synthetic negative samples using Poisson distribution modeling of SV length characteristics, filtering candidates that overlap >50% with adjacent true SVs. Our study uses all false positive calls directly from Sniffles2 output as negatives, with the intent of curating a model for identification of real-world false positives. CSV-Filter reports 95.31% F1 on HG002 / GRCh38 Sniffles2 calls, comparable to our 95.7% RF F1 across pooled datasets.

It is however important to highlight that our study is limited by leveraging only two data sets each obtained from GIAB and taking one SV caller without any filters as baseline. While these may represent the most curated SV benchmark sets, they still are limited by the number of SV, insertion and deletions types only and individuals, thus might not represent the full spectrum of genomic complexity. The somehow limited training data might very well also be a contribution to the larger models not outperforming more simplistic models overall. The relative value of deep learning models may increase in more challenging contexts, such as in cancer genomes with complex somatic rearrangements, in highly repetitive or polyploid genomes, or when analyzing variants in underrepresented populations. These arenas represent critical frontiers for future benchmarking. Furthermore, the interpretability success of sparse autoencoders points to a compelling future direction: the use of foundation models and their interpretability layers as automated feature discovery systems. The features they identify could be formalized and integrated into the next generation of simpler, more informed classifiers, creating a virtuous cycle between deep learning discovery and classical model deployment.

In summary, our systematic benchmark arrives at a clarifying and impactful conclusion: simpler, more understandable models are not merely sufficient but are fundamentally key to effective SV filtering. While the field often gravitates toward ever more complex architectures for marginal or perceived gains, we demonstrate that for the established task of germline SV filtering, this pursuit yields diminishing returns. The inherent intricacy of these advanced models often obscures their practical downsides, including substantial computational costs, reduced transparency, and a reliance on opaque features. By contrast, classical machine learning applied to well-curated genomic features provides a more practical and often superior balance of accuracy, efficiency, and interpretability. This finding reflects a critical principle for method development: sophistication should serve a clear biological or technical need, not precede it. Our results argue for a renewed focus on robust, interpretable tools as the dependable core of genomic analysis, ensuring that innovation in machine learning translates directly into reliable biological insight.

## Methods

### Dataset Preparation and Benchmarking

High-confidence SV callsets from the Genome in a Bottle (GIAB) consortium were used as ground truth for model training and evaluation [21]. For training and validation, Sniffles2 (v2.3.3, default parameters) call sets of two GIAB samples: HG002 (Ashkenazi trio son) and HG005 (Han Chinese trio son), were assessed using their corresponding benchmark sets aligned with the GRCh37 and GRCh38 reference genomes for HG002, and GRCh38 for HG005, using Oxford Nanopore Technologies (ONT) long reads [29]. Variants were benchmarked using Truvari (v4.1.0) with accompanying high-confidence BED files. Truvari parameters included size range 50bp-1Mb (--sizemin 50 --sizemax 1000000), 10% reciprocal size tolerance (--pctsize 0.10), coordinate-based matching without sequence similarity (--pctseq 0), and PASS-filter variants only (--passonly). The reported TP and FP variants were combined to have an overall dataset of 50,795 SVs, from which 44,842 are true positives (TPs) and 5,953 are false positives (FPs), distributed across three genome-reference combinations (HG002-GRCh37: 9,159 TPs/1,026 FPs; HG002-GRCh38: 17,404 TPs/3,192 FPs; HG005-GRCh38: 18,279 TPs/1,735 FPs). SAE training and evaluation focused on GRCh38-aligned datasets due to computational constraints, while feature-based models were evaluated across all available reference builds. All models were evaluated using stratified 80/20 train-test splits (40,636 training, 10,159 test) with stratification ensuring balanced representation of variant classes (TP/FP), SV types (insertions/deletions), and genome sources. This partitioning strategy was maintained consistently across all model types to enable direct performance comparisons, while leave-one-genome-out (LOGO) cross-validation was particularly used for the high-parameter vision models to ensure generalization. SAE models were trained unsupervised on all 40,437 GRCh38 variants; downstream classification using SAE features employed an independent 80/20 split.

### Feature Engineering and Selection

For each SV, 15 alignment-based features were extracted from the corresponding BAM, VCF, and reference FASTA files. These features covered size and copy number, read mapping and quality, sequence context and caller-specific attributes.

For size and copy number metrics, the base-10 logarithm of the absolute SV length was reported as log_svlen. Local coverage-based features were computed from read alignments. depth_ratio was defined as the mean per-base coverage across the variant interval divided by the mean of the 1 kb flanking regions upstream and downstream, while depth_mad represented the median absolute deviation of per-base coverage within the variant interval. Allele balance (ab) was calculated as the proportion of reads covering the SV relative to the sum of reads across the variant and flanks. cn_slop characterized larger-scale copy number context by comparing coverage within the SV to that in a distal upstream region located 5-25 kb away. Read mapping and quality features characterized local disruptions in alignment patterns. mq_drop was computed as the difference between the median mapping quality (MAPQ) of reads overlapping the SV and the flanking regions. clip_frac represented the fraction of reads containing ≥10 bp soft or hard clipping operations, while split_reads was the number of reads flagged as supplementary or containing an SA tag. read_len_med captured the median query length among reads overlapping the variant. strand_bias quantified the absolute difference between forward- and reverse-strand read fractions across the variant locus. Sequence context features were derived from the reference genome sequence extracted ±500 bp around each SV. gc_frac was defined as the proportion of G and C bases within that window, homopolymer_max as the maximum length of any sequence with identical bases, and lcr_mask as a binary indicator for low-complexity sequence, set to 1 when ≤ 2 unique nucleotides were present. Caller-specific features were drawn from the VCF INFO fields: support_read represented the number of supporting reads reported by the variant caller, and svtype_DEL was a binary indicator for deletions (SVTYPE=DEL). All features were computed per variant, with deletions evaluated between their reported start and end coordinates and insertions analyzed using a ±500 bp window centered on the insertion site. Regions with insufficient coverage or read data were assigned missing values (NaN). The resulting feature matrices were combined across datasets and written to CSV files for subsequent analysis.

For statistical comparison of true positive (TP) and false positive (FP) variant sets, distributions of each feature were evaluated using two-sided Mann-Whitney U tests. Effect sizes were computed using Cohen’s d, which measures the standardized mean difference between distributions independent of sample size and is defined as the standardized difference in means between TP and FP distributions. Features with |d| > 0.2 were interpreted as showing a meaningful effect. Effect sizes were calculated per dataset and combined using inverse-variance weighting to account for differences in sample size and precision across genome-reference combinations.

### Sparse autoencoder implementation

Genomic context around SV breakpoints was encoded using the Evo2-7B foundation model. For each SV, 1000 bp windows flanking both breakpoints were extracted from GRCh38 reference sequences and processed through Evo2 to generate 4,096-dimensional hidden-state vectors from Layer 26, previously used by the AI interpretability company Goodfire for model analysis. 1000 bp was motivated by the Evo2-7B context window of 8,192 bp. TopK sparse autoencoders were trained with a grid search across 31 configurations varying feature dimensions (2,048-32,768) and sparsity levels (k=4-128). Training employed a multi-component loss function:

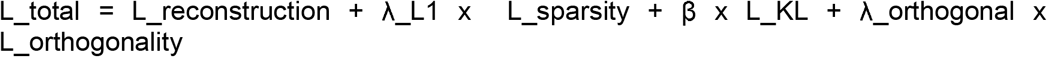

where λ_L1=0.05, β=0.5, and λ_orthogonal=0.01.

For determination of the ideal configuration, a stratified sample of 5,000 SVs was chosen containing a representative number of insertions and deletions and size categories (small <1kb, large ≥1kb), with minimum representation of 100 variants per category. Each SAE was trained for 400 epochs. The optimal configuration was then retrained on the complete HG002 and HG005 datasets aligned to GRCh38 (40,437 SVs) to generate the final SAE model used for feature extraction and evaluation. The activation patterns for the learned latent dimensions were further analyzed using odds ratios (ORs) to determine bias toward TP or FP, and cross-referenced with the 15 biologically relevant features previously mentioned.

Two complementary evaluation metrics were used to determine SAE feature quality:

- Hard F1: Direct Feature-Based Classification: For each test sample, count active FP-biased features. If ≥ threshold t, classify as negative (reject). Sweep threshold t from 1 to k_active, select optimal F1-score. This tests whether learned features can directly discriminate SVs without additional machine learning.
- Test F1: Logistic Regression Performance: Train logistic regression on SAE features on an independent 80/20 split, ensuring no data leakage in the downstream classification task. This assesses performance when SAE features are integrated into downstream ML pipelines.

Highly discriminative and interpretable atoms (those with OR > 2.0 or < 0.5 and with clear preference toward certain genomic features) were characterized through correlation analysis and manual inspection of activation patterns across diverse genomic contexts, using the University of California Santa Cruz (UCSC) Genome Browser [30].

### Image encoding

Directly following the CSV-Filter methodology [18], local read alignment evidence was converted into multi-channel images suitable for computer vision. For each candidate SV:

1. All reads were gathered overlapping the variant locus within a ±500 bp window.
2. Alignment patterns were extracted from CIGAR strings.
3. 4-channel 224×224 pixel images directly with four separate channels, where each channel captured specific CIGAR alignment operations: matches, deletions, insertions and soft-clips.
4. To enable compatibility with ResNet models, these 4-channel images were converted to 3-channel RGB format using a grayscale encoding where Match=1, Deletion=2, Insertion=3, and Soft-clip=4, then replicating this grayscale across all three RGB channels. The 4-channel to 3-channel RGB conversion maintained consistency across all vision models including ResNet and diffusion models, enabling direct performance comparison.

### Diffusion Model

Diffusion models were implemented for anomaly detection, trained exclusively on TP samples to learn patterns of genuine SVs. The standard denoising diffusion probabilistic model (DDPM) framework with forward diffusion process was employed:

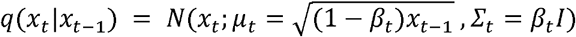

where β_t represents a noise schedule that gradually increases from small values (β_1 ≈0.0001) to larger values (β_T ≈ 0.02). Three U-Net architectures were evaluated: Small (917K parameters, 32→128 channels), Medium (14.2M parameters, 64→512 channels), and Large (56.8M parameters, 128→1024 channels). The noise schedule was optimized across standard (β: 0.0001→0.02, T=1000), low noise (β: 0.00005→0.01, T=1000), high noise (β: 0.0002→0.03, T=1000), and fast schedule (β: 0.0001→0.02, T=500) configurations. Models were trained with AdamW optimizer (learning rate=1e-4, weight decay=1e-4) using group normalization with 8 groups. Samples with a reconstruction error below a given threshold were classified as FP, with optimal thresholds determined by grid search across the 1st-99th percentile range. Because the diffusion model was trained exclusively on TP samples in an unsupervised manner (no test set labels used during training), threshold selection represents calibration of the decision boundary rather than model fitting and is an alternative to choosing a naive threshold of the median reconstruction error. This mirrors real-world deployment where practitioners would calibrate thresholds using available labeled data after unsupervised model training. Latent representations for mutual information analysis were extracted by concatenating feature maps from four encoder stages of the U-Net Large architecture: bottleneck layer (1,024 dimensions), second encoder (1,024), third encoder (512), and fourth encoder (256), yielding 2,816 total dimensions after flattening and concatenation.

### ResNet Models

Both standard ImageNet pre-trained ResNet architectures (ResNet34, ResNet50, ResNet50×2) and VICReg self-supervised models were trained. Training used frozen backbones with single trainable linear classification heads (dropout p=0.2). Data augmentation included mild color jitter while preserving alignment structure. All models were evaluated using stratified 80/20 train-test splits (40,636 training, 10,159 test) with stratification ensuring balanced representation of variant classes (TP/FP), SV types (insertions/deletions), and genome sources. This partitioning strategy was maintained consistently across all model types to enable direct performance comparisons, while leave-one-genome-out (LOGO) cross-validation was particularly used for these high-parameter vision models to ensure generalization.

Overall model performances were determined using precision, recall, F1-score, and area under the receiver operating characteristic curve (AUC). Paired t-tests across cross-validation folds were used to assess statistical significance of performance differences. For ensemble methods, 5-fold stratified cross-validation was repeated across 5 random seeds (25 total evaluations) to ensure robust performance estimates. Feature independence was quantified using mutual information analysis and canonical correlation analysis (CCA). Mann-Whitney U tests were used to identify discriminative features (p < 0.05) within each modality.

All code was executed with Python 3.13 in Jupyter Notebooks on an NVIDIA A100 GPU (40 GB VRAM) in Google Colab. The pipeline used the following packages and versions: scipy 1.14.0, numpy 2.0.0, pandas 2.2.2, matplotlib 3.9.2, scikit-learn 1.5.2, cyvcf2 0.31.1 [31], pysam 0.22.0, psutil 6.0.0, sniffles 2.3.3 [6], tqdm 4.66.5 and truvari 4.1.0 [23]. Experiments were implemented in PyTorch 2.5.1 with torchvision 0.20.1 and torchaudio 2.5.1 (CUDA 11.8 build) and transformers 4.44.2 was used for pretrained model backbones. Random seeds were fixed for all runs. The full notebooks are available in the accompanying GitHub repository for replication.

## Data availability

Oxford Nanopore Technologies long-read sequencing data for GIAB samples HG002 (Ashkenazi trio son) and HG005 (Han Chinese trio son) were obtained from the NCBI Genome in a Bottle FTP repository [29].

The code used to reproduce the results and figures presented in this study is publicly available on GitHub (https://github.com/aka133/sv_ml_filtering) and on Zenodo (https://zenodo.org/records/18277210), and is distributed under the MIT license.

## Acknowledgements

This work is supported by the NIH grant 4UH3NS132105-03.

## Contributions

FJS and AK designed the study. AK implemented different machine learning models and tested them. LFP and AK evaluated the calling performance. All authors contributed to the manuscript writing and editing.

